# Unimanual fatigue increases muscle excitation and local metabolic activity in the resting contralateral forearm

**DOI:** 10.64898/2026.06.30.735603

**Authors:** Lee J. Hinkle, Britton C. Scheuermann, Carl J. Ade, Thomas J. Barstow, Joshua C. Carr

**Affiliations:** Kansas State University, College of Health and Human Sciences, School of Health Sciences, Kinesiology, Manhattan, KS, USA

## Abstract

Intense unilateral muscle contractions evoke measurable activity within the contralateral neuroaxis, which can be detected with surface electromyographic activity in the resting homologous muscle. Physiological mirror activity (PMA), the unintentional increase in contralateral muscle excitation, has been implicated in cross-limb interactions and adaptations. Despite longstanding observations of PMA, it remains unknown whether this low-level muscle excitation influences local muscle metabolism. We addressed this question using a vascular occlusion test in 10 healthy adults. Surface electromyography and near-infrared spectroscopy-derived measures of tissue oxygen saturation and muscle oxygen consumption (mV̇O₂) were obtained from the resting left forearm during vascular occlusion at rest and during fatiguing unimanual contractions of the right hand. PMA in the contralateral resting arm was greater during unimanual fatigue than during rest (mean difference: 8.9%AA, 95% CI: 4.1 to 13.8; p = 0.002, g = 1.20). This increase was accompanied by a steeper rate of tissue oxygen desaturation (mean difference: −0.132 %·s⁻¹, 95% CI: −0.227 to −0.037; p = 0.012, g = −0.91) and greater mV̇O₂ (mean difference: 0.188 mL O₂·min⁻¹·100 g⁻¹, 95% CI: 0.057 to 0.320; p = 0.010, g = 0.94). Greater PMA was associated with both a faster rate of oxygen desaturation (r = −0.85, 95% CI: −0.96 to −0.46, p = 0.002) and greater mV̇O₂ (r = 0.78, 95% CI: 0.28 to 0.94, p = 0.008). These findings suggest that PMA is accompanied by increased local metabolic demand, consistent with a coupling between unintentional muscle excitation and oxygen extraction in the resting limb.

**New and noteworthy:** Physiological mirror activity is an established feature of intense unimanual contractions, yet its metabolic relevance has remained unknown. We show that unimanual fatigue increases contralateral muscle excitation, accelerates oxygen desaturation, and increases muscle oxygen consumption in the resting forearm during vascular occlusion. The magnitude of mirror activity was strongly associated with both oxygen desaturation and oxygen consumption, providing evidence that involuntary contralateral muscle excitation is accompanied by increased local metabolic demand in the resting limb.

## Introduction

Intense unilateral motor actions increase muscle excitation in homologous muscles of the resting contralateral limb. This response, classically termed motor irradiation (1, 2) and more recently as physiological mirror activity (3), reflects the balance of excitatory and inhibitory signals conveyed through descending motor pathways to the contralateral side (4, 5). Unilateral actions are generated within bilaterally organized cortical (6–8), subcortical (9, 10), and spinal circuitry (5, 11) and depend on excitatory and inhibitory processes across these segments to lateralize motor output while suppressing unintentional contralateral outflow (12, 13).

Physiological mirror activity can be quantified using surface electromyography (sEMG) (14) by expressing the unintentional muscle excitation recorded from the resting homologous muscle during unilateral contraction relative to that same muscle’s maximal muscle excitation, obtained during a maximal contraction of the same limb. This normalized value, termed associated sEMG activity, represents the fraction of maximal muscle excitation expressed involuntarily during a unilateral task. The magnitude of the associated sEMG activity emanating from the resting contralateral homologous muscle scales with the intensity of unilateral motor output and also rises with fatigue (15, 16). With associated sEMG activity in the resting limb reaching ∼10–40% of maximal muscle excitation, this degree of involuntary activation would be expected, in principle, to increase motor unit activity, twitch force generation, and as a result, some degree of metabolic demand. Such changes would be expected to manifest as increased oxygen consumption and reduced tissue oxygen saturation within the active musculature. However, whether this low-level involuntary excitation is sufficient to produce measurable changes in local metabolic activity remains unknown.

Therefore, the purpose of the present study was to determine whether physiological mirror activity during maximal unimanual contractions increases local muscle deoxygenation and oxygen consumption in the resting contralateral forearm. We addressed this question using a vascular occlusion test while sEMG and NIRS-derived tissue oxygen saturation (StO₂) and muscle oxygen consumption (mV̇O₂) were recorded from the resting left forearm during occlusion under two conditions: at rest and during fatiguing unimanual contractions of the right hand. Our hypothesis was that unimanual fatigue would increase associated sEMG activity (%AA), accelerate the decline in tissue saturation, and increase muscle oxygen consumption in the resting contralateral forearm compared with control conditions. Additionally, we explored whether the magnitude of associated sEMG activity was related to the reduction in tissue oxygen saturation and the increase in muscle oxygen consumption to determine whether there is a physiological coupling between involuntary contralateral muscle excitation and local muscle metabolism.

## Methods

### Experimental Design

This study employed a within-subject repeated-measures design to determine whether physiological mirror activity increases metabolic activity in the resting contralateral forearm during unimanual fatigue. Participants completed two experimental conditions involving a 3-min vascular occlusion test of the left (non-dominant) forearm with simultaneous NIRS and sEMG recordings. During the control condition, the vascular occlusion test was performed at rest.

During the unimanual fatigue condition, the vascular occlusion test was performed while participants concurrently completed fatiguing unimanual handgrip contractions with the right hand throughout the occlusion period. The order of conditions was fixed to minimize the influence of fatigue-induced carryover effects. All testing was completed during a single laboratory visit, with a 10-min washout period between conditions, consistent with previous work (17, 18). The primary outcomes were tissue oxygen saturation slope (%·s⁻¹) during the vascular occlusion test and the derived estimate of muscle oxygen consumption, both used as indices of local metabolic activity, as well as associated sEMG activity, all recorded from the resting left forearm. A graphical overview of the experimental design is shown in Figure 1.

**Figure 1.**
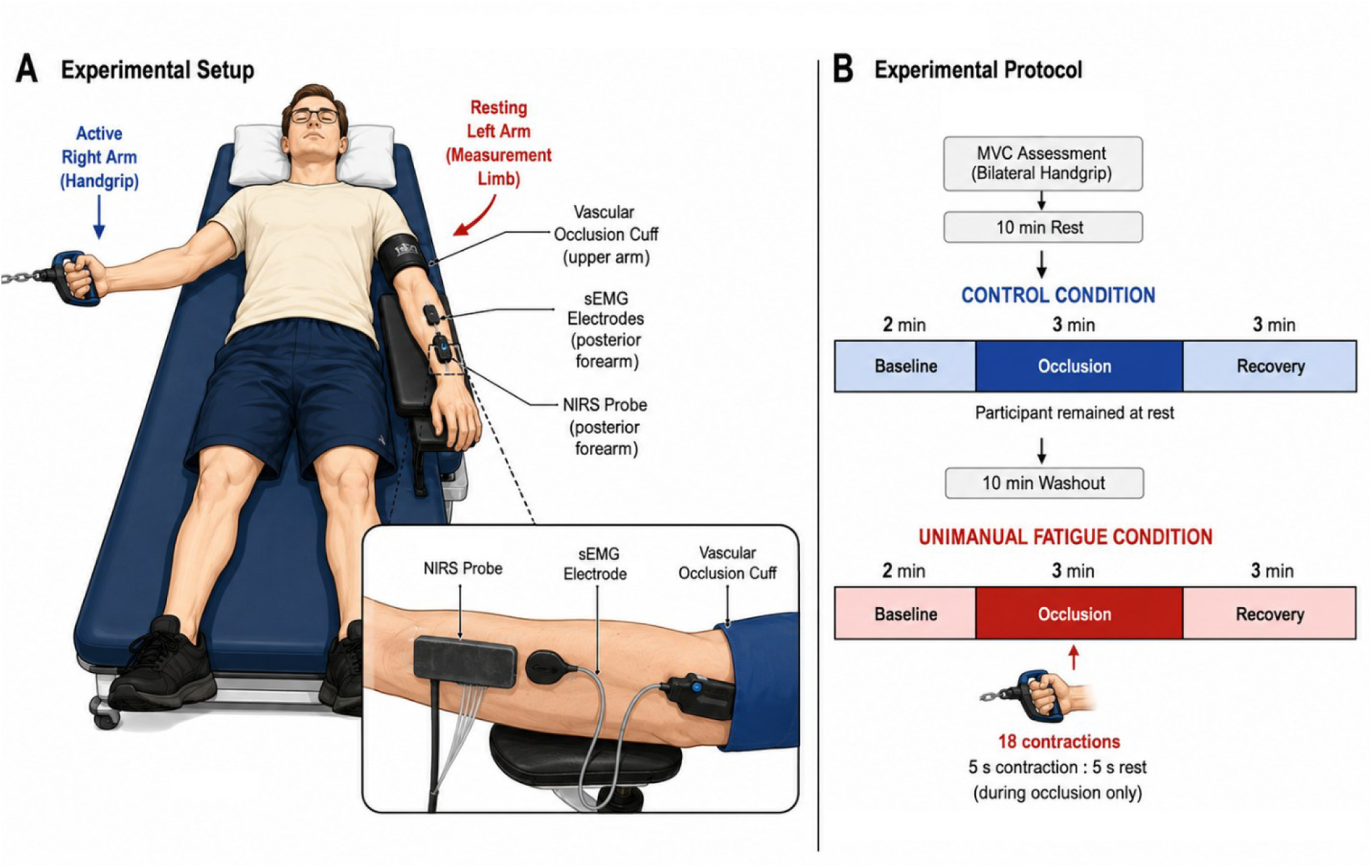
Experimental setup and study protocol. Experimental design and instrumentation used to assess physiological mirror activity and local metabolic responses in the resting contralateral forearm during unimanual fatigue. (A) Experimental setup. Participants were positioned supine on a padded treatment table. The dominant (right) hand performed intermittent maximal handgrip contractions during the fatigue condition, whereas the non-dominant (left) forearm remained at rest throughout both experimental conditions. Surface EMG sensor and the NIRS probe were positioned over the left extensor carpi radialis, and a pneumatic cuff placed around the proximal left upper arm was inflated to induce vascular occlusion. The inset illustrates the relative placement of the sEMG sensor, NIRS probe, and vascular occlusion cuff on the posterior aspect of the left forearm. (B) Experimental protocol. Following maximal voluntary contraction (MVC) assessment and a 10-min rest period, participants completed a control vascular occlusion test consisting of a 2-min baseline, 3-min vascular occlusion, and 3-min recovery while remaining at rest. After a 10-min washout period, participants completed an identical vascular occlusion protocol while simultaneously performing 18 intermittent maximal handgrip contractions (5-s contraction:5-s rest duty cycle) with the right hand during the 3-min occlusion period.

### Participants

Ten healthy adults completed the study (n = 10; 4 female, 2 hormonal contraceptives). For females, menstrual cycle phase was not controlled because testing was completed during a single laboratory visit. Participant characteristics were age, 22.2 ± 5.5 yr, and body mass index, 25.2 ± 2.9 kg·m⁻². Eligibility was determined using a medical history questionnaire. All procedures were approved by the Kansas State University Institutional Review Board (IRB-12918), and written informed consent was obtained before participation. The study adhered to the ethical principles of the Declaration of Helsinki, except for prospective registration in a publicly accessible database.

### Experimental Procedures

Experimental testing began with the assessment of maximal handgrip force. All procedures were performed with participants positioned supine on a padded treatment table. Maximal handgrip force and maximal sEMG amplitude were obtained from the left forearm, and maximal sEMG amplitude was used to normalize the associated contralateral sEMG activity, as described below. After these initial measurements, participants rested for 10 min before completing the control vascular occlusion test. Each vascular occlusion test consisted of a 2-min baseline period, a 3-min occlusion period, and a 3-min recovery period. Following a 10-min washout period, participants completed the unimanual fatigue condition using the same vascular occlusion test parameters. sEMG and NIRS from the left resting forearm was sampled continuously during the 3-min occlusion period in both conditions. The analyses focused on the wrist extensors because associated sEMG activity has been shown to be greater in the extensors than in the flexors during unimanual handgrip (19).

### Control & Fatigue Conditions

During the control condition, participants remained at rest on the treatment table while viewing a visual force feedback display set at 0% and were instructed to remain still throughout the vascular occlusion test. During the unimanual fatigue condition, participants performed intermittent maximal handgrip contractions with the right hand using a 50% duty cycle (5 seconds contraction, 5 seconds rest) throughout the vascular occlusion test, for a total of 18 contractions. Participants received visual feedback of both their maximal force reference and active force output and were instructed to contract as close to their maximal force as possible. Strong verbal encouragement was provided throughout the task.

### Maximal Voluntary Contraction and Force Measurement

Maximal voluntary contraction (MVC) force was obtained during unimanual handgrip using a modified handgrip dynamometer attached to a tension-compression load cell (Interface Inc., Scottsdale, AZ; model SSM-AJ-500), similar to previous work (19). The dominant hand was assessed first, followed by the non-dominant hand. Following a brief familiarization and warm-up, maximal handgrip force was obtained during a 5 second MVC trial with real-time visual force feedback displayed on a monitor during all testing contractions.

### Surface Electromyography

Before sensor placement, B-mode ultrasonography was used to identify the extensor carpi radialis (ECR) muscle belly of the left forearm to guide sensor placement. For most participants, the sensor was approximately 30% of the distance between the medial epicondyle of humerus and the styloid process of ulna. The skin was then prepared with isopropyl alcohol. Surface electromyography (sEMG) was recorded from the ECR using a 4-channel Galileo sensor (Delsys Inc., Natick, MA) (20) with 2.5 mm diameter EMG contact area, positioned near the centroid of the muscle. sEMG signals were sampled at 2,222 Hz and band-pass filtered at 20–450 Hz. sEMG amplitude was quantified as the root mean square (RMS) using a 250-ms moving window. RMS values were averaged across the four channels within the sensor to obtain a representative signal for the ECR. Maximal sEMG amplitude was defined as the highest 500-ms mean RMS value obtained during maximal voluntary handgrip testing, and associated sEMG activity during the experimental conditions was expressed relative to the sEMG activity during ipsilateral maximal voluntary contraction. Associated sEMG activity was calculated for each of the 18 contractions during the unimanual fatigue condition and a time-locked comparison of the same time intervals were captured from the control condition using custom R scripts to obtain 18 five second bins for both conditions. Each 5 s bin contained the average sEMG activity for that time epoch and were then averaged across bins to obtain a single representative measure for each condition. More specifically, the associated sEMG activity was quantified as the ratio of sEMG amplitude recorded from the left extensor carpi radialis during the control and right hand unimanual fatigue conditions (EMG_Resting_) to sEMG amplitude recorded from the same left forearm during left-hand maximal contractions (ipsilateral MVC). Associated sEMG activity (%AA) was calculated and expressed as a percentage:

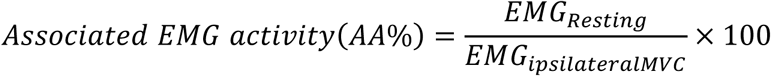

### Near-Infrared Spectroscopy, Vascular Occlusion, and Outcome Derivation

Tissue oxygen saturation (StO₂) was measured using a wireless continuous-wave near-infrared spectroscopy device (OxiplexTS; ISS Inc., Champaign, IL, USA). The probe was positioned over the left forearm extensor compartment, aligned with the extensor carpi radialis muscle belly at approximately 30% of the distance from the lateral epicondyle of the humerus to the styloid process of the radius, and placed distal to the sEMG sensor. StO₂ signals were sampled at 2 Hz. Vascular occlusion was induced via inflation of a rapidly inflatable pneumatic cuff (E20/AG101, D. E. Hokanson, Inc., Bellevue, WA, USA) and placed proximally on the upper arm to 300 mmHg. Successful occlusion was confirmed by a rapid decline in the StO₂ signal following cuff inflation. During both conditions, StO_2_ signals were low-pass filtered using a 10th-order Butterworth filter with a cutoff frequency of 0.1 Hz. Signals were visually inspected for quality, and all recordings were retained for analysis. StO₂ slope (%·s⁻¹), was calculated as the linear regression slope of StO₂ during the first 60 s of vascular occlusion using least-squares estimation (21, 22). More negative slopes indicate a faster decline in StO₂ and are interpreted as greater local oxygen extraction under ischemic conditions.

Muscle oxygen consumption (mV̇O₂; mL O₂·min⁻¹·100 g tissue⁻¹) was estimated from the initial rate of change in the [heme] difference signal ([heme]diff = oxy-[heme] − deoxy-[heme]) during the first 60 s of occlusion, quantified as the linear regression slope of [heme]diff versus time (23). Because the OxiplexTS system reports chromophore concentrations in hemoglobin-equivalent units, whereas myoglobin contributes substantially to NIRS signals in skeletal muscle, the output variables were multiplied by a factor of four to restore total [heme] units before calculation of [heme]diff and mV̇O₂ (24–26). Accordingly, the slope of [heme]diff was converted to mV̇O₂ using established assumptions regarding hemoglobin oxygen-binding capacity, tissue density, and molar gas volume (23).

### Statistical Analysis

An a priori power analysis was not performed because directly comparable data for the present experimental model and outcome measures are not available. The sample size was determined based on preliminary pilot data within our laboratory that displayed large effects for the greater StO₂ slope during unimanual fatigue. Within-subject differences between conditions were examined using two-tailed paired t-tests between conditions for all outcomes, with results reported as mean differences and corresponding 95% confidence intervals. The primary analyses evaluated differences in tissue oxygen saturation slope, muscle oxygen consumption, and associated sEMG activity between conditions. Exploratory analyses evaluated the association between associated sEMG activity and tissue oxygen saturation slope as well as muscle oxygen consumption using Pearson product-moment correlation coefficients. Effect sizes are reported as Hedges’ g. Statistical significance was set at α = 0.05. All analyses were performed in R version 4.5.1 (27).

## Results

Ten participants completed the study (n = 10; 4 female). All participants completed both experimental conditions and were included in the analyses.

### Primary Outcomes

Associated EMG activity averaged 13.4 ± 7.0 %AA during the unimanual fatigue condition and 4.5 ± 2.8 %AA during the control condition. The mean paired difference was 8.91 %AA (95% CI: 4.06 to 13.76; t(9) = 4.15, p = 0.002, g = 1.20), with all but one participant exhibiting greater associated activity during the unimanual fatigue condition (Figure 2). In parallel with these neuromuscular changes, the tissue oxygen saturation slope during the first 60 s of arterial occlusion averaged −0.258 ± 0.130 %·s⁻¹ during the unimanual fatigue condition and −0.126 ± 0.067 %·s⁻¹ during the control condition. The mean paired difference was −0.132 %·s⁻¹ (95% CI: −0.227 to −0.037; t(9) = −3.14, p = 0.012, g = −0.91), consistent with greater oxygen extraction in the resting limb during unimanual fatigue (Figure 3). To determine whether the greater rate of oxygen desaturation was accompanied by increased metabolic demand, muscle oxygen consumption (mV̇O₂) was examined. Muscle oxygen consumption in the resting contralateral extensor carpi radialis averaged 0.393 ± 0.202 mL O₂·min⁻¹·100 g⁻¹ during the unimanual fatigue condition and 0.204 ± 0.128 mL O₂·min⁻¹·100 g⁻¹ during the control condition. The mean paired difference was 0.188 (95% CI: 0.057 to 0.320; t(9) = 3.24, p = 0.010, g = 0.94), with all but one participant exhibiting greater muscle oxygen consumption during unimanual fatigue (Figure 4).

**Figure 2.**
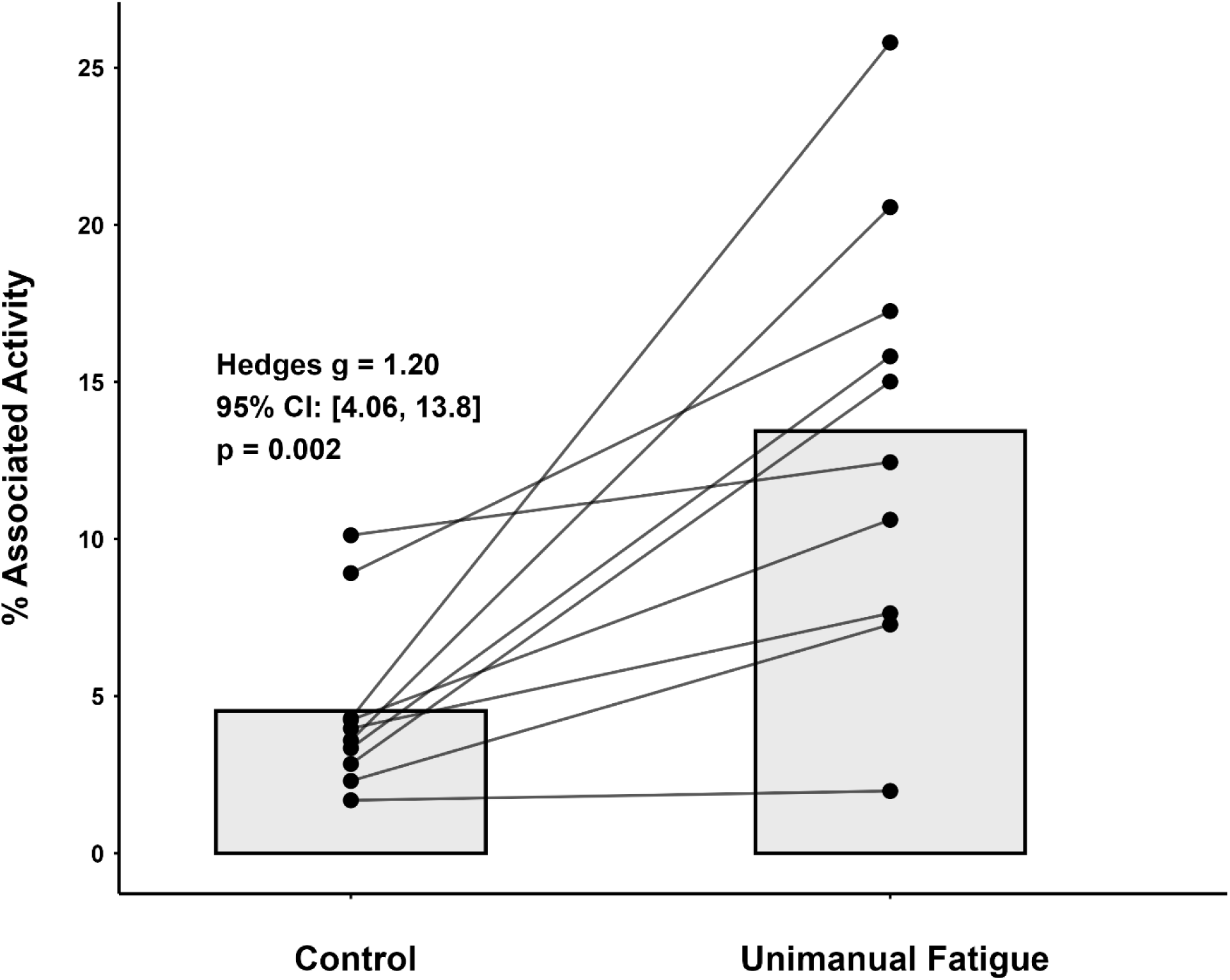
Associated sEMG activity (%AA) in the left extensor carpi radialis during control and unimanual fatigue conditions. Individual participant responses (n = 10) are shown as paired data points connected by lines, with group means displayed as bars. Associated sEMG activity was greater during the unimanual fatigue condition (13.4 ± 7.0 %AA) than during the control condition (4.5 ± 2.8 %AA), with a mean paired difference of 8.91 %AA (95% CI: 4.06 to 13.76; t(9) = 4.15, p = 0.002; g = 1.20).

**Figure 3.**
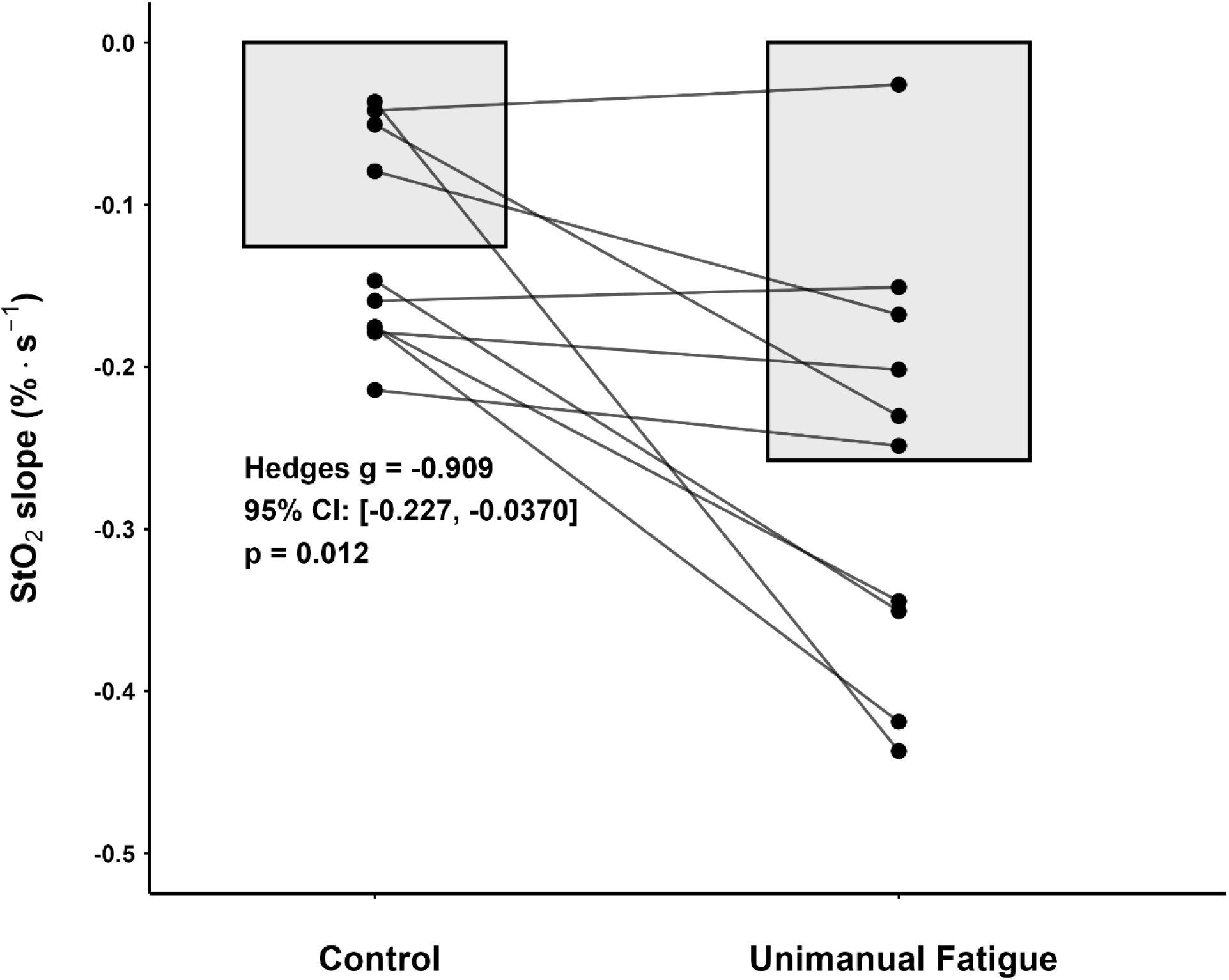
Tissue oxygen saturation (StO₂) slope (%·s⁻¹) in the left (contralateral) extensor carpi radialis during control and unimanual fatigue conditions. Individual participant responses (n = 10) are shown as paired data points connected by lines, with group means displayed as bars. The StO₂ slope was more negative during the unimanual fatigue condition (−0.258 ± 0.130 %·s⁻¹) than during the control condition (−0.126 ± 0.067 %·s⁻¹), with a mean paired difference of −0.132 %·s⁻¹ (95% CI: −0.227 to −0.037; t(9) = −3.14, p = 0.012; g = −0.91).

**Figure 4.**
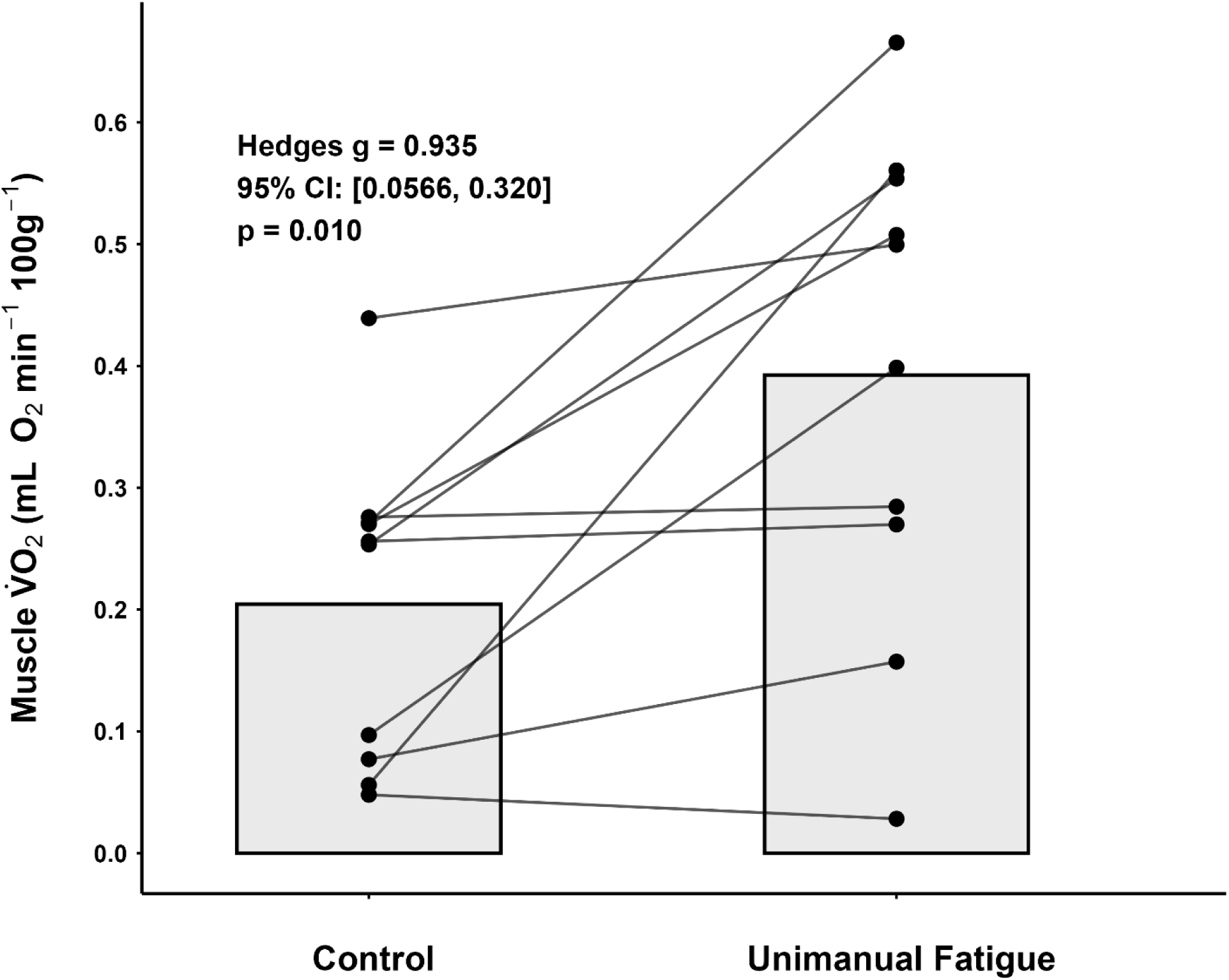
Muscle oxygen consumption (mV̇O₂; mL O₂·min⁻¹·100 g⁻¹) in the left (contralateral) extensor carpi radialis during control and unimanual fatigue conditions. Individual participant responses (n = 10) are shown as paired data points connected by lines, with group means displayed as bars. Muscle oxygen consumption was greater during the unimanual fatigue condition (0.393 ± 0.202 mL O₂·min⁻¹·100 g⁻¹) than during the control condition (0.204 ± 0.128 mL O₂·min⁻¹·100 g⁻¹), with a mean paired difference of 0.188 mL O₂·min⁻¹·100 g⁻¹ (95% CI: 0.057 to 0.320; t(9) = 3.24, p = 0.010; g = 0.94).

### Relationship Between Associated Activity and Metabolic Responses

Because associated sEMG activity, StO₂ slope, and muscle oxygen consumption differed between conditions, additional analyses were performed to determine whether the magnitude of contralateral muscle excitation was related to local metabolic responses during the unimanual fatigue condition. Associated sEMG activity was strongly correlated with StO₂ slope, such that greater contralateral excitation was accompanied by a more rapid decline in tissue oxygen saturation (r = −0.845, 95% CI: −0.962 to −0.459, p = 0.002). Associated sEMG activity was also positively correlated with muscle oxygen consumption (r = 0.775, 95% CI: 0.284 to 0.944, p = 0.008), indicating that participants exhibiting greater physiological mirror activity also demonstrated greater oxygen consumption in the resting contralateral forearm (Figure 5A-B).

**Figure 5.**
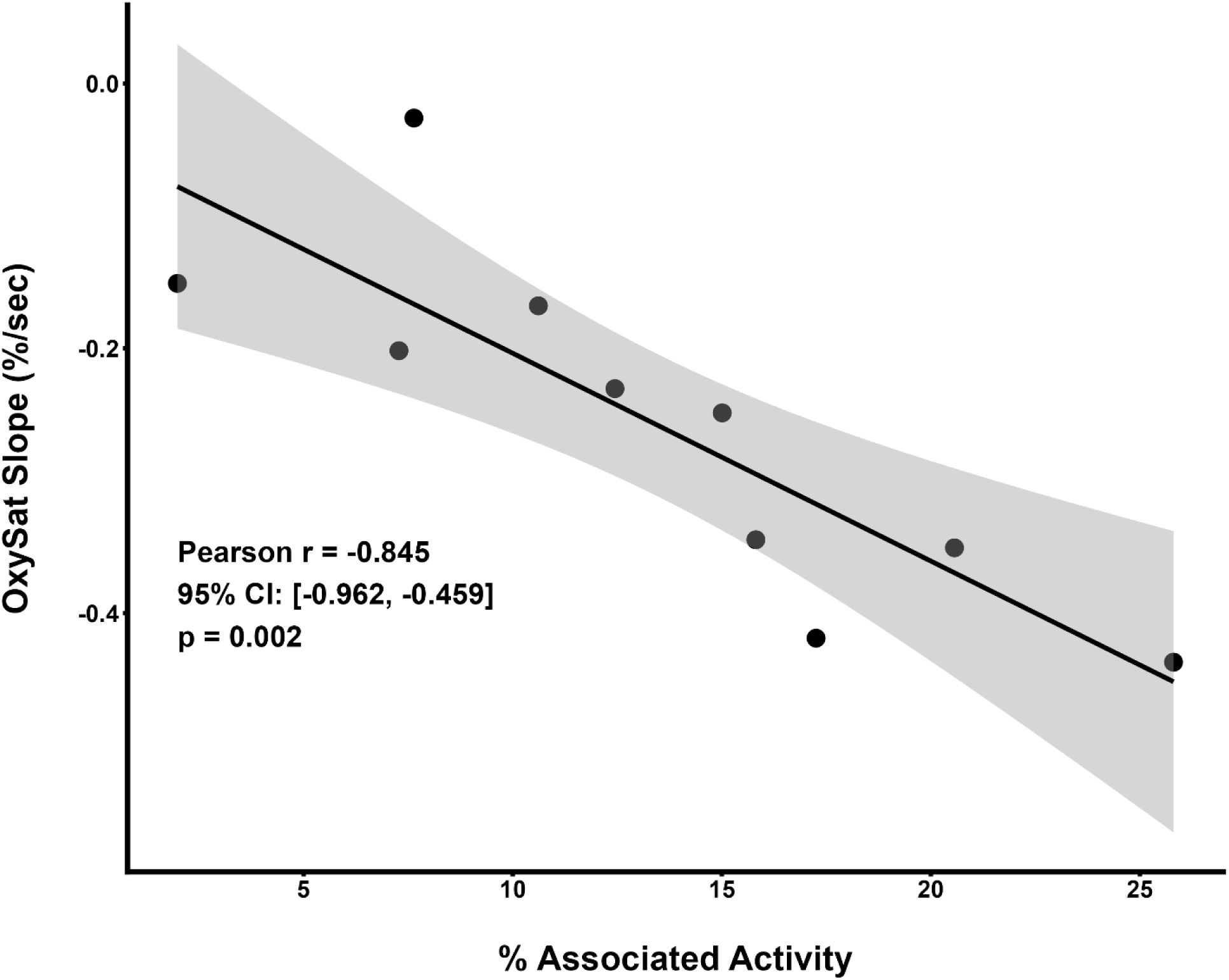
Panel A.

**Figure 5.**
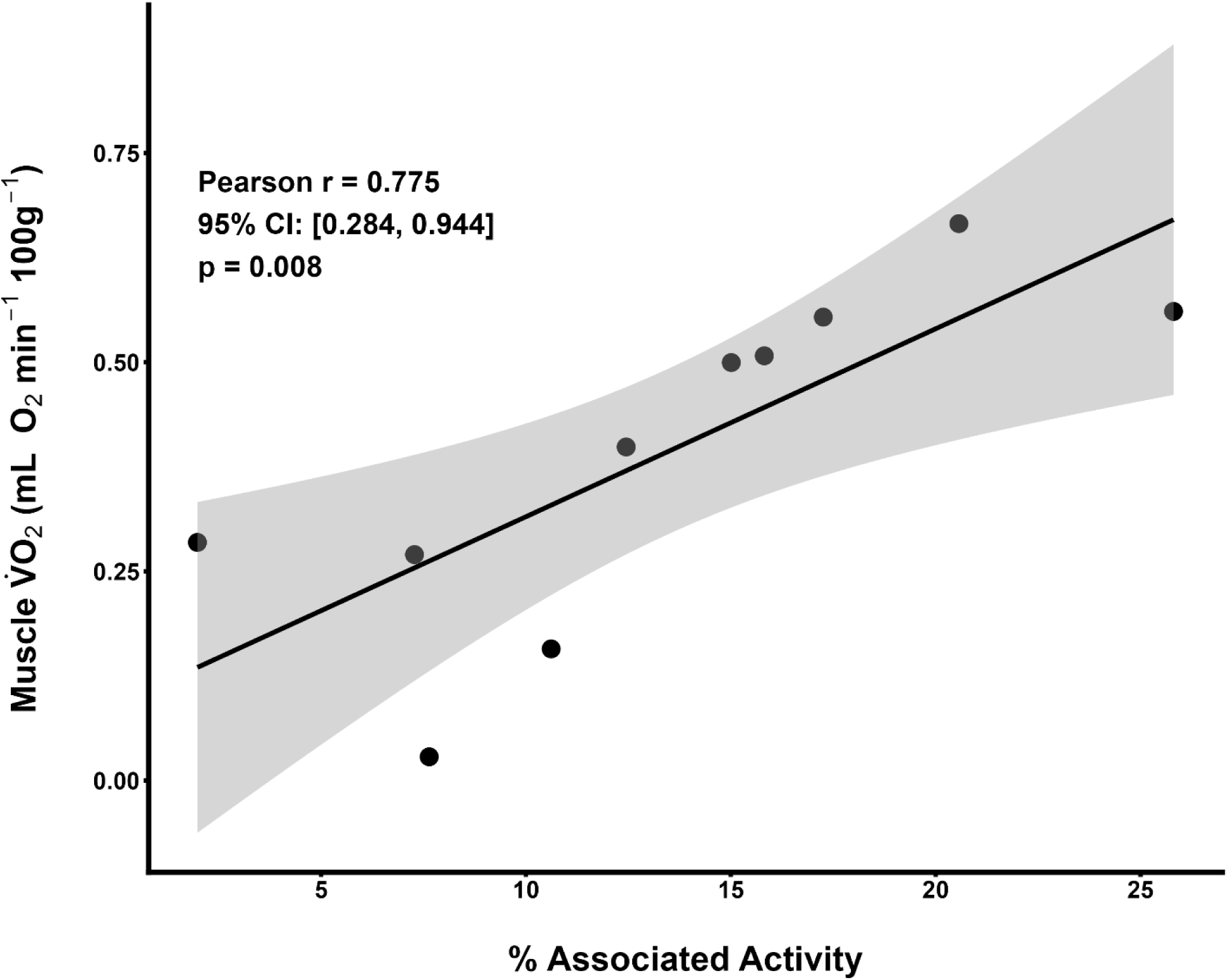
Panel B. Figure 5. Associations between physiological mirror activity and local metabolic responses during unimanual fatigue. (A) Relationship between associated sEMG activity (%AA) and StO₂ slope in the resting contralateral extensor carpi radialis. Greater associated activity was associated with a more negative StO₂ slope (r = −0.845, 95% CI: −0.962 to −0.459; p = 0.002). (B) Relationship between associated sEMG activity (%AA) and mV̇O₂. Greater associated activity was associated with higher muscle oxygen consumption (r = 0.775, 95% CI: 0.284 to 0.944; p = 0.008). Shaded regions represent 95% confidence intervals around the fitted regression lines.

## Discussion

The present study examined whether physiological mirror activity during maximal unimanual contractions is accompanied by measurable local muscle deoxygenation in the resting contralateral forearm. Using vascular occlusion while recording sEMG and NIRS-derived tissue oxygen saturation and muscle oxygen consumption, unimanual fatigue increased associated EMG activity, accelerated the decline in tissue oxygen saturation, and increased muscle oxygen consumption in the resting contralateral forearm compared with control conditions. The observed associations between physiological mirror activity, accelerated tissue oxygen desaturation, and increased muscle oxygen consumption support the interpretation that physiological mirror activity is accompanied by increased local oxygen utilization within the resting contralateral forearm. These findings extend our understanding of unimanual exercise and contralateral vascular responses (28–30) by linking unintentional muscle excitation with both accelerated deoxygenation and increased oxygen consumption within the resting forearm.

Unilateral motor tasks are generated through bilaterally organized cortical networks and rely on inhibitory mechanisms within supraspinal and corticospinal pathways to suppress overt contralateral output (7, 10, 11). During high-force or fatiguing contractions, this inhibitory control becomes progressively less effective, producing measurable excitation within homologous muscles of the contralateral resting limb. This observation has been described as physiological mirror activity (31), motor irradiation (1, 2), or contralateral muscle excitation (32) and is readily detectable using surface EMG during intense unilateral effort. Although physiological mirror activity in healthy adults rarely produces overt movement, the unintentional muscle excitation indicates activation of underlying muscle fibers and associated excitation-contraction processes.

In the active limb, increasing muscle activation is accompanied by increased oxygen utilization and extraction rate (33, 34), with metabolic demand scaling as a function of contraction intensity (35). Therefore, it is reasonable to expect that measurable involuntary excitation in the contralateral resting musculature would likewise impose a local energetic demand, even if the magnitude of activation remains substantially below that of the active limb. Previous studies demonstrated contralateral vascular and oxygenation responses during unimanual exercise (29, 30, 36), suggesting that unilateral exercise can influence the resting limb. However, because systemic circulation remained intact in these studies, changes in contralateral oxygenation could reflect centrally mediated vascular responses, altered perfusion, or broader cardiovascular adjustments rather than local metabolic activity within the resting musculature itself. Consequently, these studies demonstrated that unilateral exercise influences the resting limb but could not isolate whether physiological mirror activity was accompanied by increased local oxygen utilization. The present study sought to determine whether physiological mirror activity is accompanied by local oxygen utilization within the same contralateral tissue exhibiting involuntary excitation. By applying vascular occlusion to the resting forearm, oxygen delivery and venous washout were constrained, allowing changes in tissue oxygenation and muscle oxygen consumption to primarily reflect local oxygen extraction within the ischemic tissue volume (21, 22, 37). Under these conditions, the accelerated decline in contralateral StO₂ and the increase in mV̇O₂ observed during unimanual fatigue were more consistent with increased local metabolic demand than with systemic vascular redistribution alone. Because oxygen delivery to the forearm was constrained equally in both conditions regardless of central hemodynamic or adrenergic adjustments (21), the observed differences are most consistent with greater local oxygen extraction and oxygen consumption within the resting forearm.

We observed greater associated sEMG activity alongside steeper declines in contralateral tissue oxygen saturation and greater muscle oxygen consumption, suggesting that larger magnitudes of involuntary muscle excitation are accompanied by greater local oxygen extraction and oxygen consumption within the resting forearm. Although the present design cannot determine the precise metabolic processes responsible for this response, the most parsimonious explanation is that increased associated EMG activity reflects greater motor unit activity in the resting musculature. Even at low amplitudes, repeated motor unit discharge and low-level twitch activity would be expected to increase metabolic activity due to the contractile activity within recruited fibers. There are three lines of evidence in support of this. First, the vascular occlusion test supports this interpretation because, in the absence of blood flow, greater local oxygen utilization would be reflected as accelerated StO₂ desaturation. Second, mV̇O₂ was greater during unimanual fatigue, providing an independent estimate of increased local oxygen consumption within the resting forearm. Third, the associations between %AA and both StO₂ desaturation and mV̇O₂ further suggests that the magnitude of involuntary muscle excitation was linked to the local metabolic response. Importantly, these findings suggest that physiological mirror activity is not solely an electrophysiological observation detectable by EMG but is also accompanied by measurable metabolic consequences within the resting tissue. This extends the interpretation of physiological mirror activity beyond neural “spillover” alone and supports the notion that unintentional contralateral excitation is metabolically consequential within the resting musculature.

Physiological mirror activity has long been considered a candidate mechanism contributing to the cross-education response and prior work has attempted to determine whether this unintended excitation accounts for contralateral adaptations (38, 39). However, the evidence remains mixed as some show that mirror activity predicts cross-limb adaptations in some short-term paradigms (40), whereas others suggest that associated EMG activity is insufficient to account for contralateral strength gains (39) or the preservation of strength and muscle size during immobilization (41, 42). This dissociation suggests that mirror activity is unlikely to operate as a direct training stimulus, but it may still represent one component of a broader cross-limb response. The present findings extend this notion by showing that intense unilateral fatigue evokes a local metabolic response within the resting musculature, characterized by increased oxygen extraction and muscle oxygen consumption. Accordingly, repeated unilateral contractions may impose small but recurring neural-metabolic demands in the contralateral limb. Over time, this energetic stress and repeated metabolic activation could plausibly interact with centrally mediated motor adaptations and local cross-limb molecular signaling to contribute to adaptive processes within the untrained musculature (43). This possibility is consistent with preclinical evidence that unilateral treatments can evoke contralateral skeletal muscle responses, including immune-cell signaling (44) along with enhanced tissue remodeling and protein turnover in the opposite limb following mechanotherapy (45), and immune- and stress-response transcriptional pathways following unilateral electrical stimulation (46). This speculation provides a physiological framework by which chronic unilateral training could influence the untrained limb through the coupling of central motor drive, local contralateral oxygen utilization, and cross-limb molecular signaling. Defining these mechanisms in unilateral training paradigms will further advance our understanding of adaptive cross-limb responses.

### Methodological considerations

We note several considerations when interpreting the present findings. First, NIRS does not directly measure metabolism, but rather detects changes in the absorption characteristics of heme-containing chromophores within the sampled tissue volume. Accordingly both tissue oxygen saturation (StO₂) and the derived estimate of muscle oxygen consumption (mV̇O₂) should be interpreted as indirect indices of local oxygen utilization rather than direct measurements of metabolic flux. Further, NIRS-derived signals represent mixed tissue contributions that may include skin, adipose tissue, microvasculature, and skeletal muscle.

Therefore, we interpret the StO₂ slope primarily as an indirect estimate of local oxygen extraction and and mV̇O₂ as an estimate of local aerobic metabolic rate derived from NIRS-based changes in heme oxygenation during vascular occlusion (23, 37). The use of vascular occlusion was an intentional methodological approach designed to constrain oxygen delivery and minimize systemic blood flow influences, thereby improving interpretation of local oxygen extraction within the resting forearm. However, even under occlusion, both StO₂ slope and mV̇O₂ remain NIRS-derived estimates of local oxygen utilization rather than direct measures of tissue metabolism. The fixed-order experimental design minimized fatigue carryover effects between conditions (17, 22) but does not fully exclude potential order effects. In addition, the modest sample size should be considered when interpreting the breadth of these findings, although the within-subject consistency, large effect sizes, and relatively narrow confidence intervals support the robustness of the observed responses. Despite these considerations, the combined use of vascular occlusion, sEMG, and NIRS provides convergent evidence supporting an association between physiological mirror activity and increased local oxygen extraction and oxygen consumption in the resting forearm.

## Conclusion

We show that physiological mirror activity during unimanual fatigue is accompanied by accelerated deoxygenation and increased muscle oxygen consumption in the resting contralateral forearm, consistent with enhanced local metabolic activity. Importantly, the increase of unintentional muscle excitation (%AA) was associated with a steeper decline in StO₂ and greater mV̇O₂. These strong associations suggests that physiological mirror activity—an established feature of intense unilateral muscle contraction—produces measurable metabolic consequences in the contralateral homologous pair experiencing involuntary excitation in an intensity-dependent manner. These findings extend our understanding of cross-limb interactions by demonstrating that unimanual fatigue evokes coupled neural and metabolic responses in the resting limb, providing new insight into bimanual coupling and potential adaptive mechanisms involved in the cross-education response.

## Data availability

Data are stored on a public repository with access through the corresponding author.

## Ethics Approval

This study received ethical approval from Kansas State University’s Institutional Review Board for Human Subjects Research (ID #12918), and all subjects signed an Informed Consent document.

## Author contributions

Conceptualization: BCS, CJA, TJB, JCC; Methodology: LJH, BCS, CJA, TJB, JCC; Formal analysis and investigation: LJH, BCS, JCC; Writing – original draft preparation: LJH, JCC; Writing – review and editing: LJH, BCS, CJA, TJB, JCC; Resources: BCS, CJA, TJB, JCC; Supervision: JCC. All authors reviewed, edited, and approved the final version for submission.

